# Identification of genes that differentiate *Mannheimia haemolytica* genotypes 1 and 2 using a pangenome approach

**DOI:** 10.1101/2025.05.13.653884

**Authors:** Darien Deschner, Janet E. Hill

## Abstract

*Mannheimia haemolytica* is an opportunistic bacterial pathogen associated with the economically costly bovine respiratory disease. Two genotypes have been described, of which genotype 2 is more strongly associated with disease. Several previous studies have investigated the genomic differences between the genotypes and/or the major serotypes (1, 2 and 6) of *M. haemolytica*, however we still lack a clear basis for the greater disease association of genotype 2 (serotypes 1 and 6) and demonstrations of phenotypic differences are scarce. This work builds upon previous investigations to identify genes that differentiate the two genotypes with a particular focus on genes that may play a role in virulence and fitness in the respiratory tract microbiome. We identified 422 genotype differentiating genes in a collection of 206 unique *M. haemolytica* genomes (61 genotype 1, 145 genotype 2). Genotype differentiating genes included genotype-associated variants of a TonB-dependent siderophore receptor homolog, transferrin binding protein B, leukotoxin A, and IgA1 proteases. We also identified a genotype 1 associated lytic transglycosylase, and a genotype 2 specific highly immunogenic outer membrane lipoprotein. Genotype 2 genomes were significantly larger in size and contained more predicted protein coding genes than genotype 1 genomes. These results expand our knowledge of what differentiates the genotypes 1 and 2 of *M. haemolytica* and provides information that can be used as the basis for laboratory investigations of corresponding phenotypic differences.

## Introduction

*Mannheimia haemolytica* is a gram-negative opportunist bacterial pathogen associated with bovine respiratory disease (BRD) [1,2]. The beef industry is estimated to lose more than $3 billion per year globally due to BRD [3,4]. Two genotypes of *M. haemolytica* have been described based on whole genome sequencing and single-nucleotide polymorphism (SNP) analysis and multiple serotypes have also been defined [5,6]. The three most common serotypes (1, 2 &, 6) have been found to correspond to genotypes with serotypes 1 & 6 corresponding to genotype 2 (G2) and serotype 2 corresponding to genotype 1 (G1) [1,5,7,8]. Genotype 2 has been more frequently associated with clinical cases of BRD, however the genetic and phenotypic basis for this apparent association is unclear [1,9–14]. In addition to virulence factors such as leukotoxin, lipopolysaccharide, adhesins and proteases [11,15–17], adaptations that confer increased fitness, such as greater ability to acquire iron and other nutrients, could contribute to disease association by allowing a pathogenic strain to outcompete non-pathogenic strain(s) present in the same environment [18].

Reports of specific phenotypic differences between *M. haemolytica* G1 and G2 are scarce. G2 strains corresponding to serotype 1 have been demonstrated to invade differentiated bovine bronchial epithelial cells *in vitro* while G1 strains (serotype 2) could not [19]. The only other phenotypic differences described to date are distinct matrix-assisted laser desorption/ionization - time of flight mass spectrometry (MALDI-TOF) profiles and colony morphology [20,21].

Genotypic differences between G1 and G2 have been investigated previously using whole genome sequences. A pangenome analysis of 69 *M. haemolytica* genomes identified 112 G1 specific, and 179 G2 specific genes. This study was focused on outer membrane protein encoding genes and identified seven G2 specific predicted outer membrane protein encoding genes that were hypothesized to play a role in the greater disease association of G2 [22]. Another study examined 11 *M. haemolytica* genomes for potential differences in virulence factors between the serotypes 1, 2 and 6 [23]. The authors reported more prophages in serotypes 1 and 6 than in serotype 2, which could influence virulence due to the presence of virulence factors in these prophages. Self-targeting CRISPR spacer sequences were detected in serotypes 1 and 6, which may enhance immune evasion or adhesion during infection by modulating sialic acid expression on the cells surface. ARG-carrying ICEs were more prevalent in serotypes 1 and 6 than serotype 2, which could enhance survival of antibiotic exposure [23].

The objective of our current study was to take advantage of the greater number of *M. haemolytica* whole genome sequences now available to identify genetic differences between the two genotypes. In our examination of >200 *M. haemolytica* genomes, we gave particular attention to genes encoding proteins that may contribute to the reported differences in disease association such as those involved in virulence and/or fitness in the bovine respiratory tract. We designed an approach that allowed the detection of both genes with genotype associated presence/absence patterns and sequence variants of genes conserved in both genotypes. This contrasts with previous studies that have largely focused on the presence/absence of genes within the genotypes. Our results showed that G2 genomes are significantly larger in size and contained more predicted protein coding genes than genotype 1 genomes. We identified a total of 422 genotype differentiating genes, including several whose predicted functions may partially explain the differences in observed disease association and that have not to our knowledge been reported previously. This list of genes will inform future research to demonstrate phenotypic differences between G1 and G2.

## Materials & Methods

### Genome sequences

*M. haemolytica* whole genome sequences defined as complete by NCBI were downloaded from the NCBI Genome database on March 25, 2024, using the NCBI Datasets command line tools [24]. NCBI defines “complete” as all chromosomes are gapless and contain runs of nine or less ambiguous bases, no unplaced or unlocalized scaffolds, and all expected chromosomes present. To avoid duplication all .gca files that had a matching .gcf file were removed. To identify G1 and G2 genomes, the sequences were genotyped using an *in silico* PCR assay in Unipro UGENe v45.1 and primers from a genotype specific LAMP assay with default parameters (Mismatches = 3, Min Perfect Match =15, Max Product Size = 5000, Use ambiguous bases = True, Extract annotations = Inner, and Temperature settings = Primer 3) [25,26]. Primers from the LAMP assay included a pair targeting G1 specific adhesin pseudogene B1, G2 adhesin G (CP017538.1 locus tag: BG586_06285; Forward: 5’-GCA CAT TAA AAT ATA GCA GCT TTG-3’; Reverse: 5’-CTA AGC CAG AGT GAT CCG-3’), and leukotoxin D (*lktD*) (CP005972.1 locus tag: F382_07400; Forward: 5’-ACT GAA AA TTC ACA CTA TAG GTG-3’; Reverse: 5’-GCT AAT ATG TTT AAT TCG ACC AGT T-3’). The *lktD* primers were used to confirm identity of genomes as *M. haemolytica*. In initial testing G1 specific primers failed to produce any predicted products in our G1 reference strain and so were discarded. New G1 specific primers targeting the G1 specific adhesin pseudogene B1 were developed using Primer 3 v 4.1.0 (CP017495.1 locus tag BG548_01640; Forward: 5’-ACT AAT CTG AAG AGC GGC GT-3’; Reverse: 5’-TTG GTG GTT GCT GCT TGT AC-3’) [27–29]. G1 and G2 NCBI genomes were combined with a set of G1 and G2 genome sequences previously generated by our lab (BioProject: PRJNA1088094)(30) and reference genomes (G1 = CP017518; G2 = CP017519) for both genotypes.

A phylogenetic tree was constructed using MASH with the ani command in PanTools to verify genotype designations [30].

### Pangenome construction, annotation, and classification

A pangenome was constructed using PanTools v 4.3.2 with FASTA files, gff3 annotation files, and genotype designation as inputs [30–32]. Genomes were assigned a new unique identifier based on their input order during pangenome construction (S1 Table). Genome metrics such as genome size, GC content, number of genes, etc. were determined using the metrics command.

Pantools groups protein sequences from the input genomes into homology groups based on their similarity. Homology groups also contain the nucleotide sequences encoding the constituent protein sequence(s). Homology groups were generated using the group command with the relaxation parameter set to 2. The relaxation parameter is a combination of four sub-parameters (intersection rate, similarity threshold, mcl inflation, and contrast) where a lower number represents a stricter set of thresholds. We elected to utilize a relaxation parameter of 2 to allow for detection of minor variations that might be expected when examining sub-populations of the same species and avoid grouping of more divergent genes as might occur with a looser parameter.

Homology groups were classified by PanTools as core (present in ≥ 90% of all genomes), unique (present in only one single genome), or accessory (present in >1 genome and <90% of all genomes). Based on their prevalence within each genotype, homology groups were classified as either shared (present in ≥ 90% genomes of one genotype, regardless of their prevalence in the other genotype), exclusive (solely present in genomes of one genotype), or specific (present in ≥ 90% genomes of one genotype, and no genomes of the other genotype) [30]. To capture homology groups strongly associated with one genotype that did not meet the strict PanTools definition of specific we used the classification “genotype associated” for homology groups that were present in ≥ 95% of one genotype while also being present in > 0% and ≤ 5% of the other genotype.

Genomes were functionally annotated using InterProScan v 5.64-96.0 with SignalP v 4.1 database integrated [33–35]. Biosynthetic gene cluster annotations were generated using antiSMASH 7.1.0 with default parameters [36]. InterPro and antiSMASH annotations were added to the pangenome, with InterProScan annotations including InterPro, Pfam, GO, and SignalP annotations [30].

Multiple sequence alignments of constituent nucleotide sequences were generated for all homology groups in the pangenome with MAFFT and consensus sequences from the alignments were generated to represent the genotype specific and associated homology groups [30,37]. Consensus sequences were used as queries for BLASTx searches using the bacterial/archaeal translation code against the NCBI refseq_select database [38–42]. All other parameters were kept as default. Searches were performed March 20-25, 2025. The top hit for each search was manually recorded to provide further evidence of the predicted function, facilitate human interpretation, and identify functions that may not have been annotated by any of the other approaches. The entire pangenome database is available in S1 Dataset [37].

The nucleotide sequences of several previously characterized genes upregulated in response to iron limitation, and previously identified proteases were used as BLAST queries against the pangenome using the blast command in PanTools to determine if these previously characterized genes were present in our dataset [15,30,43]. Both alignment threshold and minimum identity were set to 90. Query nucleotide sequences are available S2 Table.

### Tet(H) gene abundance

Tet(H) genes were used as a proxy for the presence of integrative and conjugative elements (ICE) due to the strong correlation between presence of tet(H) and presence of ICE [12,44]. The reference tet(H) nucleotide sequence (Accession Number: Y15510.1 (nucleotides 2099-3301)) was downloaded from the comprehensive antibiotic resistance database (CARD) on July 26, 2024, and used as nucleotide BLAST query against the pangenome database using the inbuilt PanTools blast command. Minimum identity and alignment threshold were both set to 90%.

### Leukotoxin operons

Reference nucleotide sequences for each of the component genes of the *M. haemolytica* leukotoxin operon (*lktCABD*) were downloaded from GenBank (AF314508.2) and used as queries against the pangenome BLAST database with percent identity and alignment threshold both set to 90%. Consensus nucleotide sequences of G1 and G2 versions of all leukotoxin operon genes were extracted and translated in Geneious 2024.0.5. Predicted protein sequences for the G1 and G2 versions of all leukotoxin operon genes were aligned against one another in Geneious 2024.0.5.

### Annotation of carbohydrate active enzymes

Carbohydrate active enzymes (CAZymes) were annotated using a local installation of dbCAN v 4.1.1 software with integrated SignalP v 4.1 database and DIAMOND, HMMER, and HMMER dbcan_sub prediction algorithms [45]. Annotation was performed on each genome sequence individually using a custom BASH script [37]. dbCAN annotations that had supporting annotations from at least three tools and contained a predicted signal peptide were selected for further analysis.

### Protein structure prediction and visualization

Protein sequences were submitted to the AlphaFold Server for structural prediction [46]. Models with the highest ranking score were downloaded and visualized in PyMol version 2.5.2 (Schrödinger, LLC).

### Statistics

Welch’s t-test for unpaired data was used to compare G1 and G2 genome characteristics with *P* ≤ 0.05 considered significant and 95% confidence intervals for the difference in means were reported. A one-way ANOVA (Kruskal-Wallis) with Dunn’s test for multiple comparisons was used to determine if there was a difference in genome size between G1 and G2 with or without ICE. *P* ≤ 0.05 was considered significant. These analyses were performed in GraphPad PRISM v 10.4.1.

A Fisher’s exact test and odds ratio comparison was performed to determine if the prevalence of tet(H) was different between genotypes. Both tests were performed in Microsoft Excel for Microsoft 365 using Real Statistics Resource Pack. *P* ≤ 0.05 was considered significant.

## Results & Discussion

### Description of genome collection

*M. haemolytica* genomes designated as complete were retrieved from NCBI and after removal of duplicates, 121 genomes were available for further analysis. *In silico* PCR with modified primers from a previously published genotyping assay resulted in the expected amplicons for 108/121 genomes. No *in silico* genotyping PCR products were obtained for 13 genomes and so they were removed from further analyses leaving 108 genomes.

Three of the 13 isolates that could not be genotyped also failed to produce any predicted product for the *lktD in silico* PCR. Two of these genomes have since been removed by the RefSeq curation team for either containing too many frameshift mutations (GCF_900474405) or failing to pass a completeness check (GCF_023516295). All three genome sequences that failed to produce an *lktD* product were from sheep isolates (NCTC 10643, NCTC 10609) or from culture collections from the 1960s (ATCC 29701). Of the other ten genomes that were not genotyped by *in silico* PCR, four were from a German study examining macrolide resistance in *M. haemolytica* [44], and three were from sheep isolates collected in 1951 or 1968 and sequenced as part of the United States Department of Agriculture (USDA) *M. haemolytica* sequencing project (BioProject: PRJNA172854). The last three isolates were all generated from non-nasopharyngeal cattle samples as part of either the USDA sequencing project (GCF_000819525, GCF_023516255) or a German study (GCF_020971705) and had a CheckM completeness <92.5%.

The *in silico* PCR assay we used was selected for identification of G1 and G2 genomes as defined by Clawson et al. [5]. The original LAMP assay from which we selected primers for this process was developed using *M. haemolytica* isolates from North American cattle [25]. Thus, the most likely explanation for genomes to be un-typable by this approach (besides sequence quality issues) is that they represent genotypes other than G1 or G2.

The NCBI genomes were combined with 99 genomes from our previous work (BioProject Accession PRJNA1088094) and reference genomes for G1 and G2 as described in the Methods. One additional genome was removed following discovery that it was likely a chimera, leaving a final dataset 208 *M. haemolytica* whole genomes with 62 G1 and 146 G2 representatives (S1 Table). The reference genomes for G1 and G2 were also represented in the collection of genomes downloaded from NCBI resulting in the collection representing 206 unique *M. haemolytica* genomes. Most genomes were from isolates originating from feedlot cattle in either Canada or the United States between 2003 and 2021. Consistent with the *in silico* PCR identifications, a pangenome derived phylogenetic tree showed two distinct clades corresponding to isolates classified as G1 or G2 (Figure S1).

### Genome properties & Tet(H) abundance

The two genotypes were compared to one another on a broad set of genome metrics, such as genome size, GC content, etc., to determine if there were any high-level differences in the genomic structure and content of the two genotypes. G2 genomes were larger, had more predicted protein-coding sequences, and had slightly lower GC content than G1 genomes. No differences were identified in the number of tRNA or rRNA genes or the proportion of the genome covered by genes (Table 1).

**Table 1.**
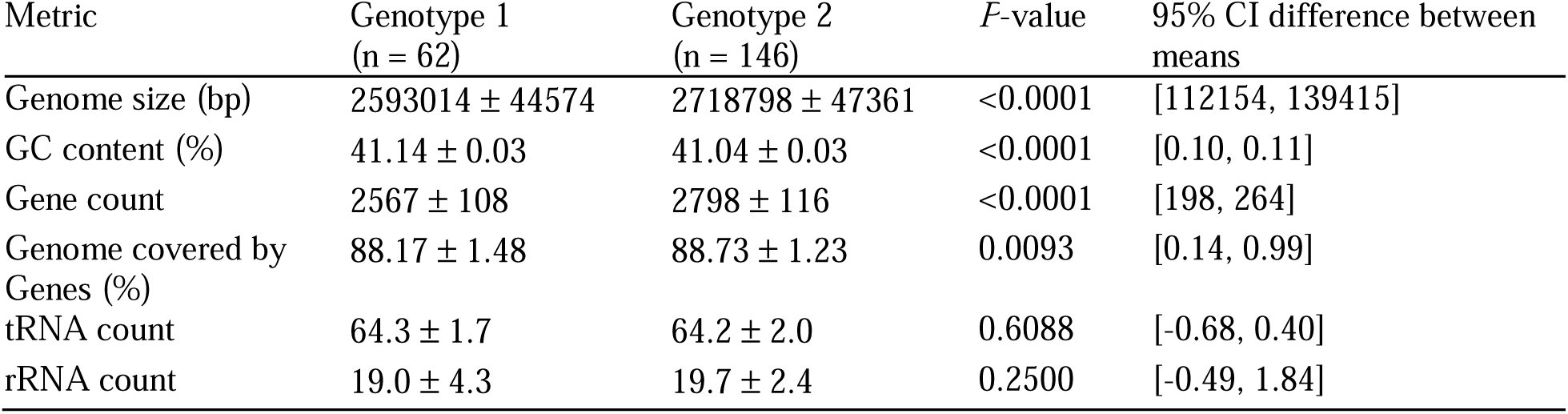
Comparison of genome features of G1 and G2. Values are mean ± standard deviation. *P*-values are from a Welch’s t-test with *P*<0.05 considered significant.

| Metric | Genotype 1<br>(n = 62) | Genotype 2<br>(n = 146) | *P*-value | 95% CI difference between means |
| --- | --- | --- | --- | --- |
| Genome size (bp) | 2593014 ± 44574 | 2718798 ± 47361 | <0.0001 | [112154, 139415] |
| GC content (%) | 41.14 ± 0.03 | 41.04 ± 0.03 | <0.0001 | [0.10, 0.11] |
| Gene count | 2567 ± 108 | 2798 ± 116 | <0.0001 | [198, 264] |
| Genome covered by Genes (%) | 88.17 ± 1.48 | 88.73 ± 1.23 | 0.0093 | [0.14, 0.99] |
| tRNA count | 64.3 ± 1.7 | 64.2 ± 2.0 | 0.6088 | [-0.68, 0.40] |
| rRNA count | 19.0 ± 4.3 | 19.7 ± 2.4 | 0.2500 | [-0.49, 1.84] |

We hypothesized that the larger size of G2 genomes was attributable to a greater prevalence of integrative and conjugative elements (ICEs), large (∼20 kb to >500 kb) mobile genetic elements that are known to be associated with antimicrobial resistance and are more frequently identified in G2 genomes [5,47]. To test this hypothesis, we used the proxy measure of the presence of tetracycline resistance gene tet(H) to determine the prevalence of ICEs due to the strong association of tet(H) with ICEs [12,44].

Tet(H) was identified significantly more frequently in G2 (67/146, 45.9%) than G1 (1/62, 1.6%) (Fisher’s *p*-value = 4.39 x 10^-12^ Odds Ratio: 0.02 ± 1.02), however, G2 genomes were larger than G1 regardless of the presence of ICE (Figure 1). The larger size and higher gene count of G2 genomes regardless of presence of ICE (Table 1) could indicate that G2 genomes carry a larger repertoire of genes that may allow them to be more adaptable to a wider variety of environments than the smaller G1 genomes. Considering average bacterial gene sizes, the difference in the median genome size of G2 without ICE and G1 (∼100 kb) could represent dozens of G2-specific genes. Additionally, the greater prevalence of ICEs in G2 suggests that this more disease associated genotype is also more likely to possess antimicrobial resistance genes.

**Figure 1.**
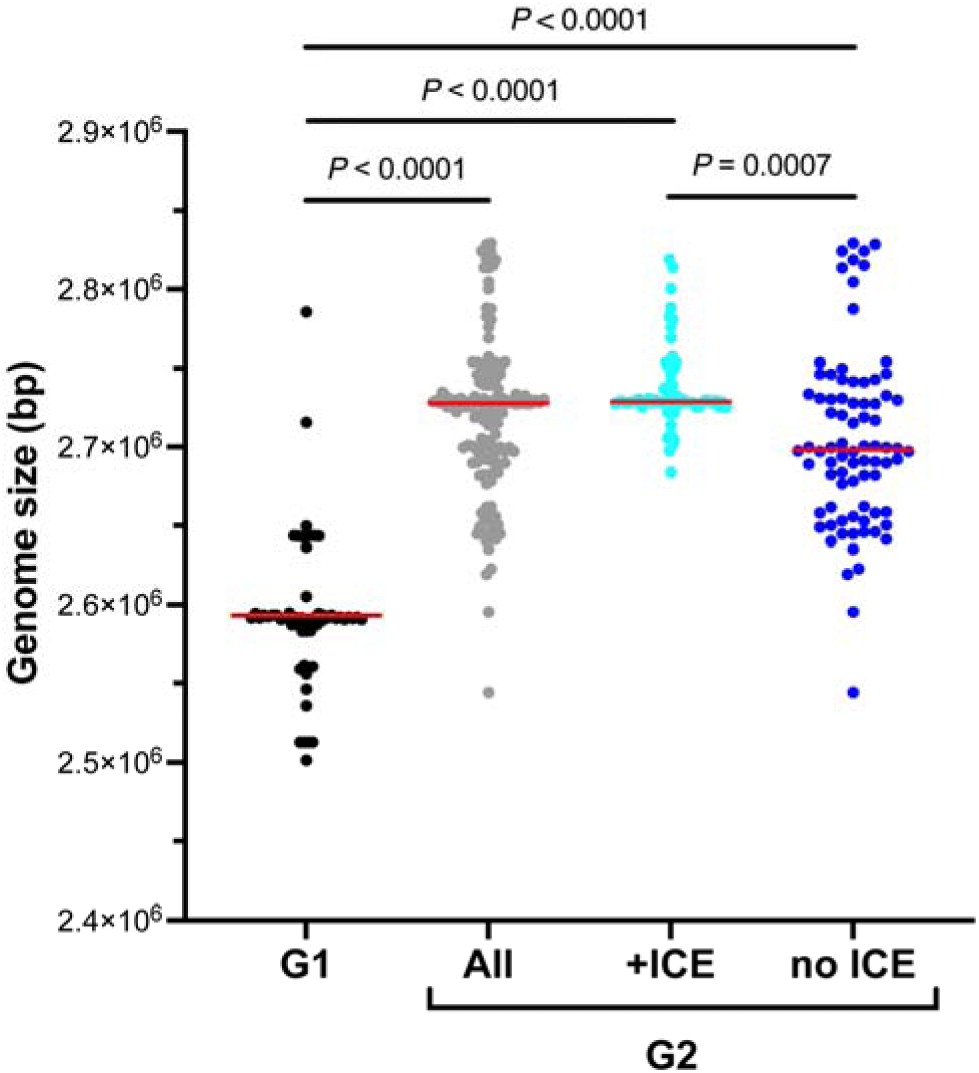
Genome sizes of Genotype 1 (G1, n = 62), Genotype 2 (G2, n = 146), G2 possessing integrative and conjugative elements (ICEs, n = 66), and G2 without ICEs (no ICE, n = 80). One way ANOVA (Kruskal-Wallis) followed by Dunn’s multiple comparisons test (corrected *P* values are shown). Red bars indicate mean.

### Pangenome analysis

The pangenome calculation resulted in the identification of “homology groups”: clusters of sequences from input genomes that exceed a given similarity threshold. We then examined the distribution of homology groups across all *M. haemolytica* genomes to identify those that are core, unique, accessory, shared, genotype specific or genotype associated as defined in the methods. A total of 4,038 homology groups were generated of which 2,007 (49.7%) were core, 1,473 (36.5%) were accessory, and 558 (13.8%) were unique (Pangenome_Classification in S3 Table).

For the current study, our interest was homology groups that differentiate the genotypes: those that are found in the large majority of representatives of one genotype and in none, or a small minority, of the other genotype. PanTools identified 135 G1 specific and 233 G2 specific homology groups. The PanTools specific category definition is strict, however, requiring a homology group to be present in >90% of one genotype and zero of the other. To identify homology groups that do not meet this strict criterion but are nevertheless strongly associated with one genotype, we examined accessory groups not initially classified as specific for those that were present in ≥ 95% of the one genotype and > 0% and ≤ 5% of the other genotype. These were classified as G1 associated (n = 21) or G2 associated (n = 34) (S3 and S4 Tables). Identification of genotype specific and associated genes within this relatively large genome collection reinforces previous findings that the two genotypes carry distinct sets of genes [22]. The pangenome results also provide evidence of the internal coherence of the genotypes, since when considered separately, G1 and G2 genomes included 2,169 and 2,342 “genotype-core” homology groups (present in >90% of genomes of the genotype), respectively.

To provide an overview of the potential phenotypic differences between the two genotypes, all specific and associated homology groups were assigned to one of nine general functional categories based on functional annotations and the results of manual BLASTx searches against the NCBI refseq_select database (Figure 2, S4 Table, S5 Table). For both genotypes, the largest number of specific and associated homology groups were classified as MGE proteins (transposases, toxin-antitoxin plasmid maintenance systems, prophage associated proteins, etc.) or hypothetical. The larger number of G2 specific and associated MGE proteins and antimicrobial resistance genes is consistent with greater ICE prevalence in G2 [9,12,22,23,44]. The large numbers of genotype specific and associated hypothetical proteins in both genotypes is also unsurprising as ∼22 % of all domain families in Pfam v32.0 are defined as “domains of unknown function” and a recent examination of >30,000 bacterial genomes and metagenomes showed that 40-60% of genes are of unknown function [48,49]. Homology groups predicted to encode unknown/hypothetical proteins, and MGE proteins were excluded from further analysis in this study but certainly represent an opportunity for future investigation.

**Figure 2.**
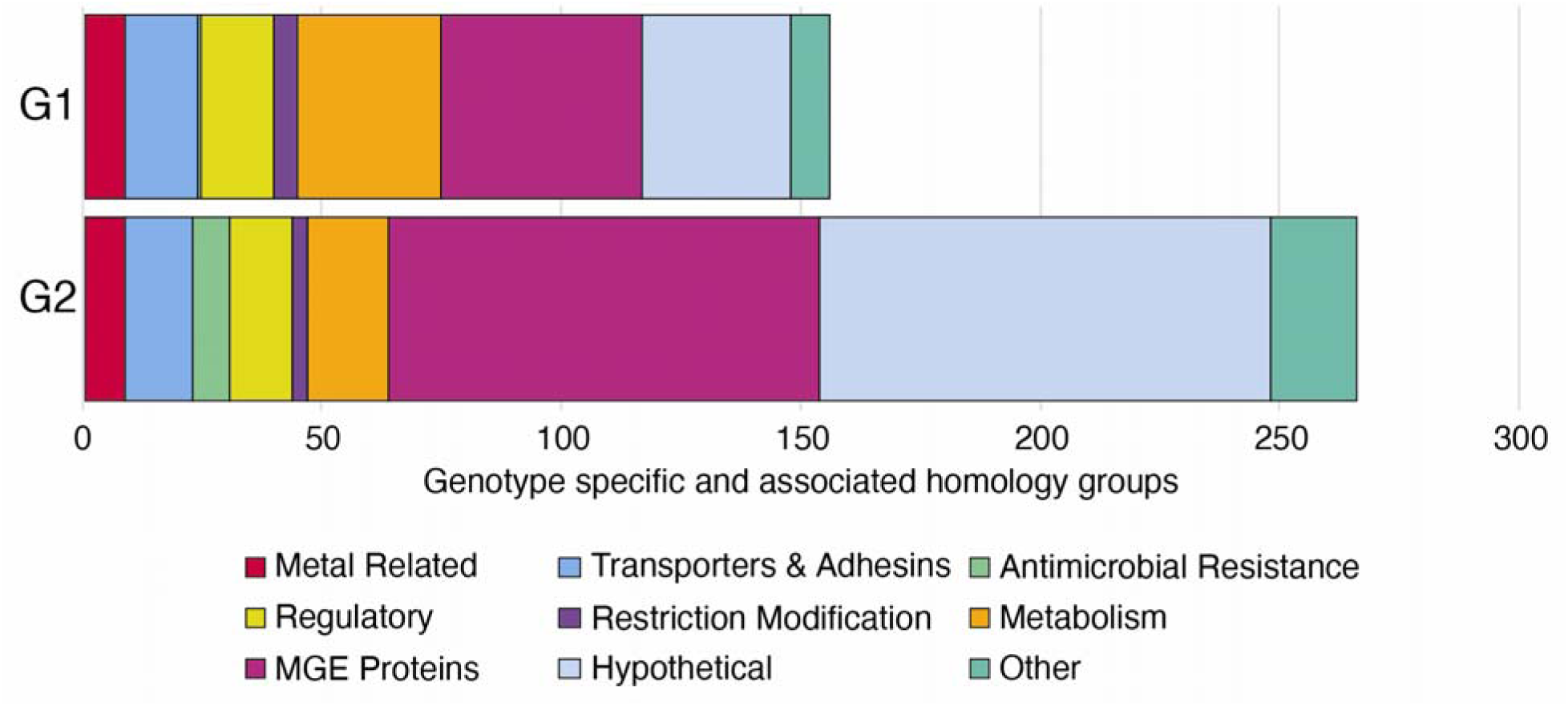
Predicted functional categories of genotype specific/associated homology groups (n = 156 for G1; 266 for G2). G1 = Genotype 1, G2 = Genotype 2. Specific groups are those found in ≥90% of the genomes of the given genotype and no genomes of the other genotype. Associated groups are present in ≥95% of the genomes of one genotype > 0% and ≤5% of the genomes of the other genotype. MGE proteins include transposases, toxin-antitoxin plasmid maintenance systems, prophage associated proteins, etc. Hypothetical proteins include those annotated as hypothetical or domain of unknown function.

Based on inspection of the complete inventory of pangenome homology groups (Table S3, S4), we identified a subset of specific/associated homology groups with functional annotations that may be associated with reported differences between disease association of G1 and G2 (Table 2). These genome features were related to iron acquisition, host immune evasion, immunogenicity and carbohydrate modification.

**Table 2.**
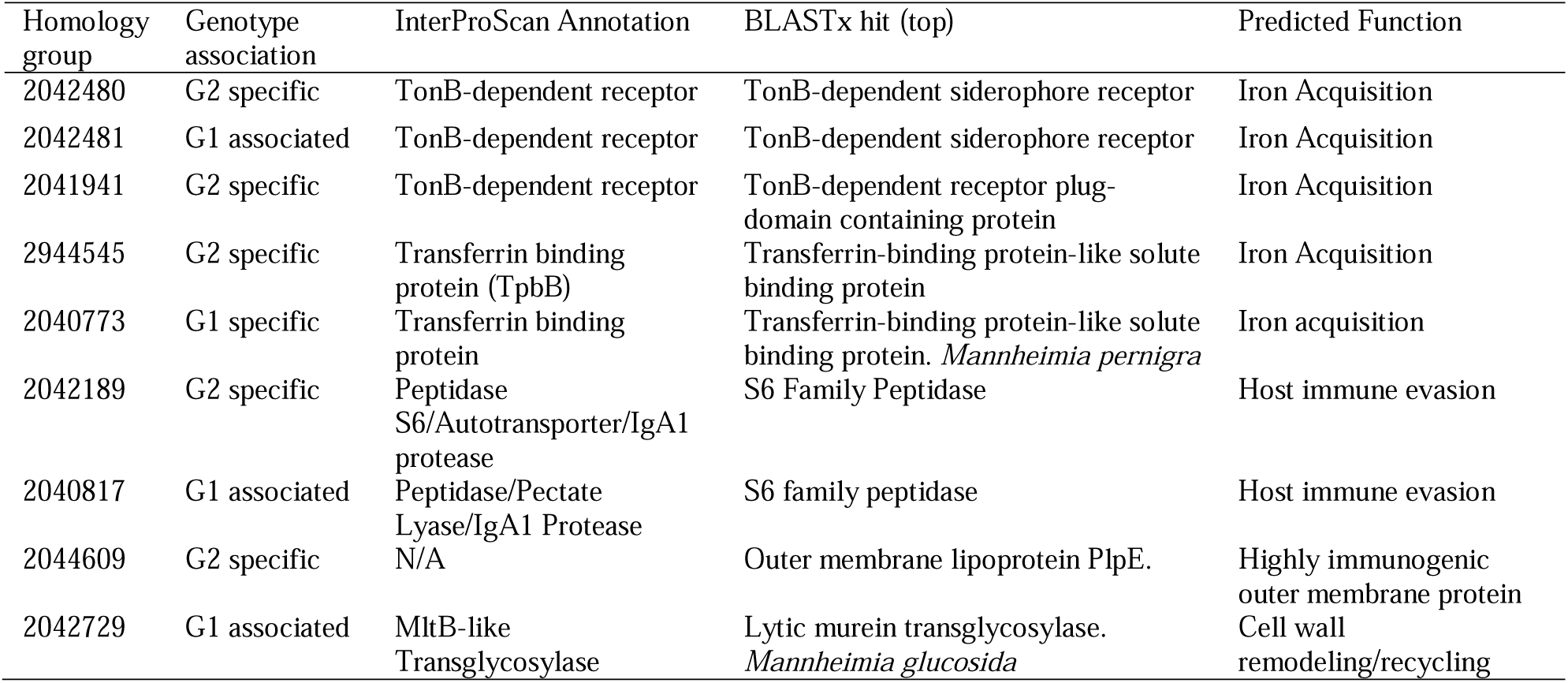
Genotype association and predicted functions of selected homology groups. Annotation notes summarize the functional annotations from InterProScan for each homology group. BLASTx search used consensus nucleotide sequence of the homology group as query against the refseq_select database. Reported top BLASTx hits were from *M. haemolytica* unless otherwise stated.

### Iron acquisition

Iron is essential for bacterial growth and bacterial pathogens must compete with each other and with the host for the scant iron typically available in their environment [50,51]. Animal pathogens like *M. haemolytica* are faced with concentrations of available iron well below their needs, which has necessitated the evolution of mechanisms for acquiring iron including siderophores, transferrin binding proteins, and haem binding proteins [52]. Given the essential nature of iron, and the generally iron limited conditions of the respiratory tract, the ability to acquire iron more readily than other bacteria provides a considerable advantage. The identification of several genotype associated, and specific homology groups potentially involved in iron acquisition was of interest since these differences may be associated with fitness in the respiratory tract and thus with disease association (Table 2).

Homology groups 2042480 (G2 specific) and 2042481 (G1 associated) were both annotated as encoding a TonB-dependent siderophore receptor. TonB dependent receptors (TBDTs) are transporters often involved in iron acquisition, specifically in the uptake of small, secreted iron scavenging molecules known as siderophores [53,54]. Homology groups 2042480 and 2042481 also produced BLAST hits to a previously characterized siderophore (ferric enterobactin) receptor (*frpB*) homolog that was upregulated in *M. haemolytica* in response to iron limitation [43]. The consensus amino acid sequences of these homology groups were 84.4% identical, and both contained consensus TonB-dependent transporter (TBDT) motifs. Predicted 3D structures demonstrated the canonical 22 β-strand barrel structure with an N-terminal plug domain characteristic of TBDTs (AlphaFold ranking_score = 0.95 for both) (Figure 3A). The amino acid sequence differences between the G1 and G2 sequences were concentrated in the extracellular loops responsible for substrate binding, suggesting that these two proteins may differ in their ability to bind their substrate [53,54] or they may bind different substrates (Figure 3B). Identification of a putative siderophore receptor in *M. haemolytica* is unexpected as *M. haemolytica* is not known to produce or utilize siderophores and has been demonstrated to be unable to utilize ferric enterobactin so it is likely that the identified TBDTs have some other function besides siderophore uptake [11,43].

**Figure 3.**
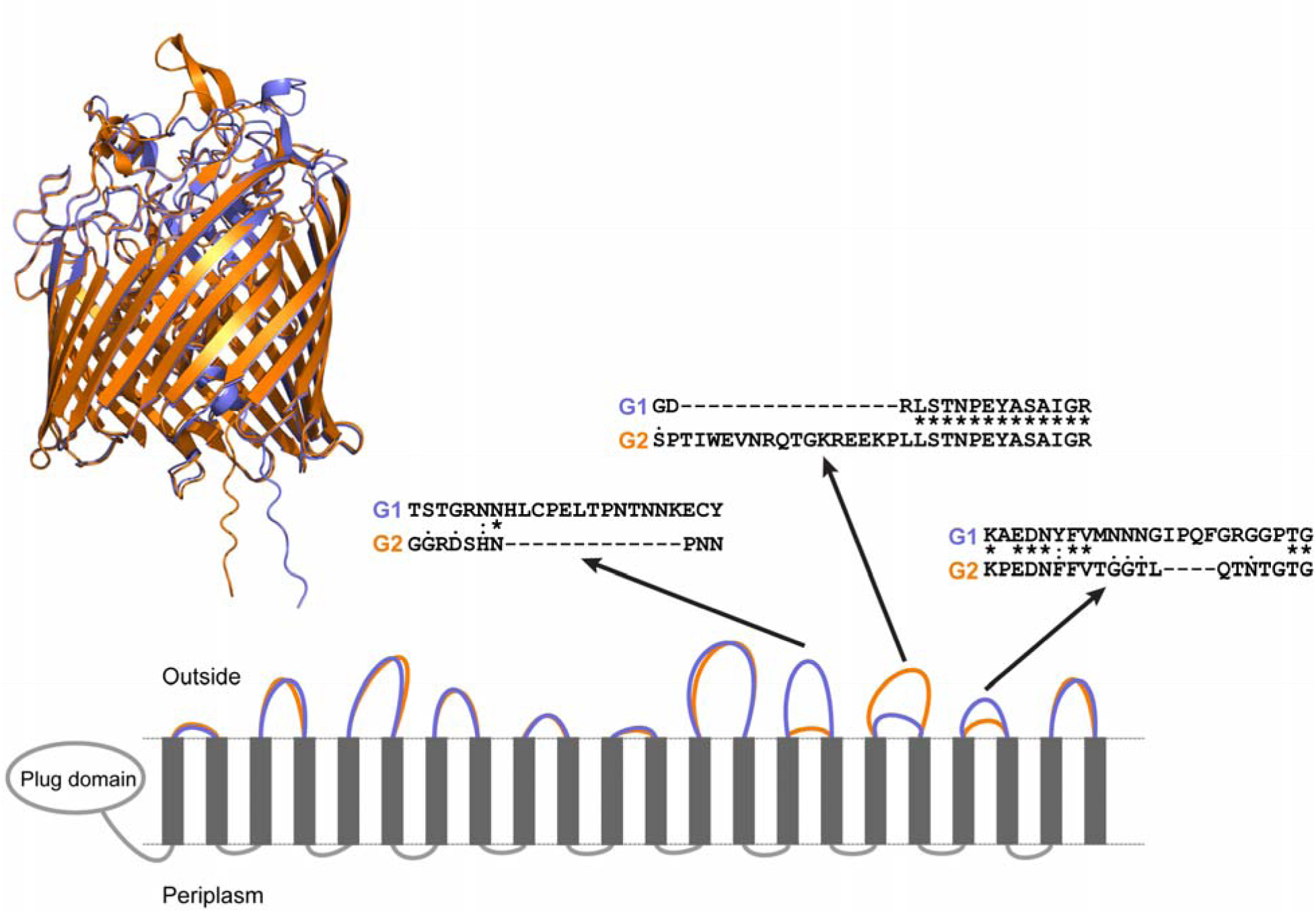
A) Predicted 3D structures of Genotype 1 (G1, Purple) & Genotype 2 (G2, Orange) ferric enterobactin receptor (frpB) homolog proteins. 3D structure conforms to canonical TonB-dependent receptor structures. B) 2D representation of tertiary structure of G1 and G2 frpB homologs. Each grey rectangle represents one of the 22 β-strand sheets, and the size of the extracellular/periplasmic loops are shown. Differences in the amino acid sequence for select extracellular loops are shown in further detail. Difference in extracellular loops may contribute to differences in substrate binding efficiency and/or result in interactions with completely different substrates. G1 AlphaFold ranking_score = 0.95; G2 AlphaFold ranking_score = 0.95.

G2 specific homology group 2041941 was also predicted to encode a TBDT. No similar G1 specific/associated protein was identified and the consensus sequence of 2041941 did not correspond to previously identified genes upregulated in response to iron limitation [43]. The identification of another G2 specific TBDT suggests that G2 may have a greater ability to uptake iron and/or other substrates. TBDT have been implicated in transport of a variety of substrates in gram-negative bacteria including siderophores as well as vitamin B_12_ and carbohydrates [54].

Further examination is required to determine the function of this TBDT and whether it provides a competitive advantage to G2 over G1 strains. Other iron acquisition proteins include transferrin binding proteins A and B (*tpbAB*), both of which are found in *M. haemolytica* [11]. Transferrin is an iron-carrying glycoprotein found in mammalian circulatory systems that transports iron throughout the body and serves as a sink for any excess free iron released into the circulatory system [55]. TpbB is a surface lipoprotein that extends outwards from the outer membrane to the extracellular space where it binds to transferrin and brings the transferrin closer to TbpA, a TBDT in the outer membrane that transports the transferrin into the periplasm [55]. To investigate any genotype associations of transferrin binding proteins, previously characterized *tbpA* and *tbpB* nucleotide sequences from *M. haemolytica* were used as BLAST queries against the pangenome database. Core homology group 2044546 contained hits for *tbpA* while *tbpB* hits were solely identified in G2 specific homology group 2044545 with no hits in any G1 genomes. These results suggest that *tbpA* is highly conserved within *M. haemolytica* and likely plays a key role in iron metabolism. In contrast, while no BLAST hits were produced to the query *tbpB* nucleotide sequence in G1 genomes, G1 specific homology group 2040773 was predicted to encode for a transferrin binding protein. The amino acid sequences of G1 and G2 variants of the putative TbpB proteins (homology groups 2040773 and 2044545) were compared and found to be 56.5% identical and 81.2% similar. Despite the relatively low similarity of these two proteins, the predicted 3D structures were very similar to one another as well as the crystal structure for the TbpB from *Actinobacillus pleuropneumoniae* (G1 ranking_score = 0.88, G2 ranking_score = 0.93, Figure 4). A difference in the TbpB structure between the genotypes was observed in the N lobe with G2 appearing to have a longer alpha-helix structure in the N-lobe handle region than G1 or *A. pleuropneumoniae*. This region of the protein contains residues important for binding of iron-loaded transferrin [56]. Genotype variation of a transferrin binding protein further suggests they may differ in their ability to acquire iron, which may partially explain observed differences in disease association between the two genotypes. The genotype variants of TbpB identified here may also reflect host preferences of the genotypes as TbpB is selective for host transferrin and differences between TbpB proteins has been partially attributed to this host selectivity [55].

**Figure 4.**
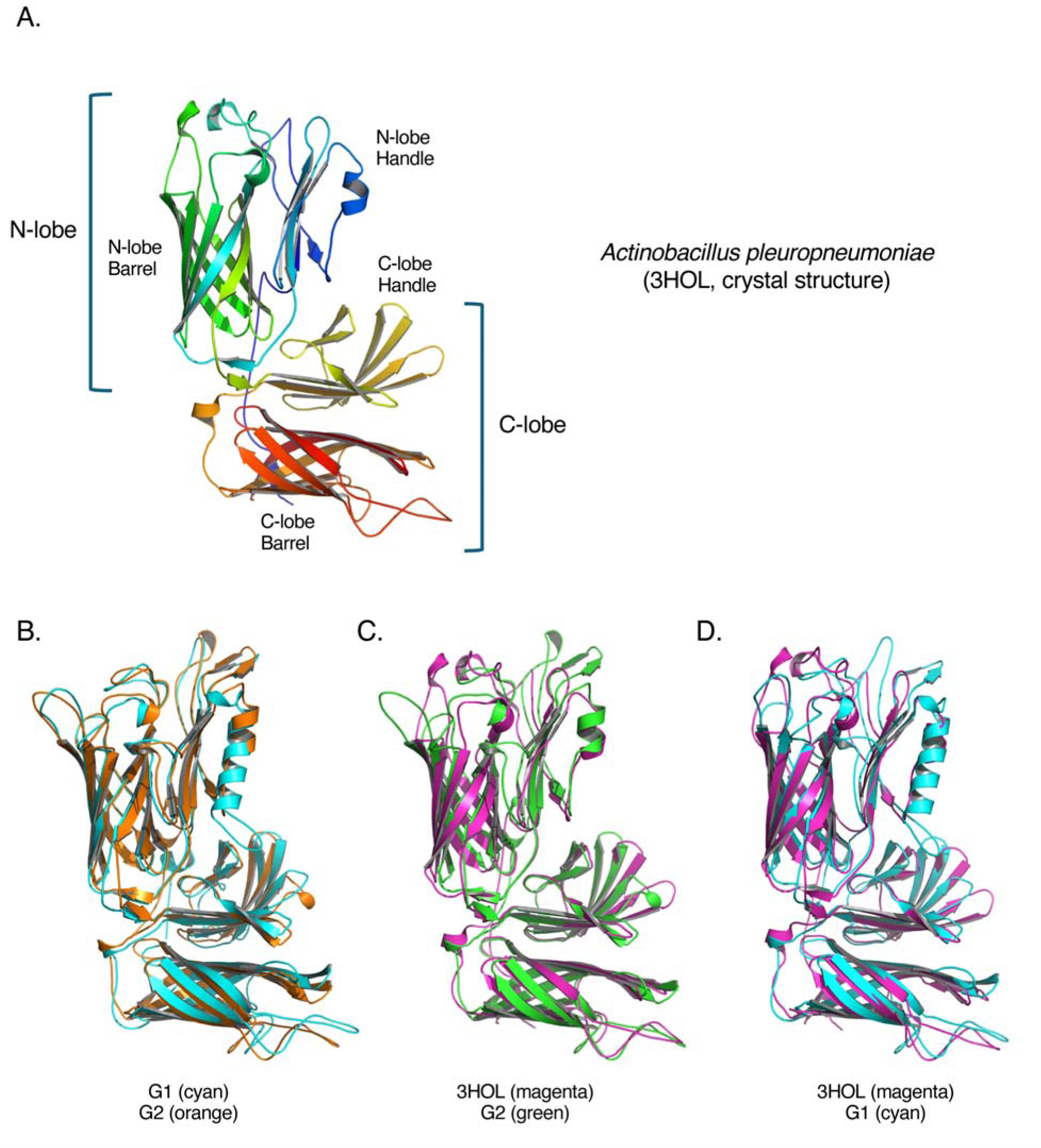
Predicted structures of G1 and G2 variants of a predicted transferrin binding protein B (TbpB). (A) The crystal structure of TbpB from *Actinobacillus pleuropneumoniae* (PDB accession 3HOL [56]) was used as a reference. Major domains include the N-lobe and C-lobe, each containing a beta-barrel and a “handle”. The reference structure is rainbow coloured from N terminus (blue) to C terminus (red). (B) Alignment of G1 and G2 variants. (C) G2 aligned to reference. (D) G1 aligned to reference. The G1 variant has an elongated alpha-helix in the N-lobe handle domain relative to G2 and the reference. (G1 ranking_score = 0.88, G2 ranking_score = 0.93).

### IgA protease

IgA proteases are metalloproteases that cleave immunoglobulin A (IgA), a critical component of the mucosal immune system. Two IgA proteases (IgA1 and IgA2) have been previously identified and characterized in a G2 (serotype 1) *M. haemolytica* isolate [15]. We identified a G2 specific IgA protease (homology group 2042189) and a G1 associated IgA protease (2040817) (Table 2). To determine the relationship of these sequences to the previously reported IgA proteases, a BLAST search was performed using previously reported nucleotide sequences for IgA1 and IgA2 as queries against the pangenome database (IgA1:NZ_DS264709 Locus Tag: MHA_RS00165IgA2: NZ_DS264612.1 Locus Tag: MHA_RS09765). At least one BLAST hit was returned for IgA2 in all but five of the included genomes with hits grouped into five homology groups (2041035, 2041036, 2041399, 2041400, and 2043723). Homology groups 2041305 and 2041306 were accessory, while 2041399 and 2041400 were unique suggesting some minor variations are likely present within the pangenome for this IgA2 protease. Homology group 2043723, however, was a core group suggesting that this IgA2 protease is a core gene of *M. haemolytica* and is not genotype differentiating. In contrast, a BLAST hit was returned for IgA1 in 146/209 of the genomes with hits grouped into one of three homology groups (2040817, 2041298, and 2042189). Homology group 2041298 was found in only one genome so was not examined any further. Group 2040817 is a G1 associated group while 2042189 is a G2 specific group and thus these groups appear to represent genotype variants of an IgA protease.

Comparison of the consensus sequences of the two IgA1-like homology groups revealed that the nucleotide sequence of 2042189 (G2 specific) is 4,512 bp consistent with previous descriptions of IgA1 in G2 *M. haemolytica* [15], while the sequence of 2040817 (G1 associated) was only 2,281 bp. An alignment of the two variants showed that they were 94% identical over 2,281 shared positions and that the G1 associated sequence is a truncated version of the other. The *M. haemolytica* IgA1 protease contains three functional domains: a large, N-terminal peptidase domain, pertactin-like passenger domain and a C-terminal autotransporter domain [15]. The product of the truncated G1 associated gene would include the peptidase domain, but lack part of the pertactin-like passenger domain and the autotransporter, and thus may not encode a complete or functional IgA1 protease.

The observation of a G2 specific IgA1 sequence and possibility of genotype variation remains of interest and warrants further investigation since differences in IgA protease activity could relate to difference(s) in their ability to evade the host immune system and thus may contribute to differences in disease association.

### Highly immunogenic outer member lipoprotein

*Pasteurella* lipoproteins (Plp) are a group of several immunogenic outer membrane lipoproteins identified in *M. haemolytica* [57] and PlpE has been investigated as a potential vaccine target for *M. haemolytica* because of its immunogenicity [58–60]. Aside from the immunogenic role of PlpE relatively little is known regarding its function although it is important as a vaccine target and thus clinically and diagnostically significant.

G2 specific homology group 2044609 was predicted to encode PlpE, however no corresponding G1 homology group was identified. Previous investigations of PlpE sequence diversity across serotypes showed that the serotype 1 and 6 (G2) sequences were highly conserved whereas serotype 2 (G1) PlpE sequences were extremely heterogenous, however, antibodies raised against serotype 1 PlpE cross reacted with serotype 2 bacteria [61]. To identify potential G1 variants of the G2 *plpE*, a reference *plpE* gene (NZ_CP023044.1, c780551-779481, locus tag: CKG22_RS03975) was used in a BLASTx search with relaxed parameters. When percent identity threshold was reduced to 70% and alignment threshold to 50% a hit to G1 specific homology group 2042820 was observed.

The identification of a sequence encoding a G2 specific PlpE and possible G1 variant suggests that the genotypes may differ in immunogenicity, however further research is needed to investigate the prevalence and immunogenicity of Plp variants across *M. haemolytica* genotypes.

### Leukotoxin operon

Leukotoxin (Lkt) is a critical virulence factor in *M. haemolytica* and is produced by both genotypes (all serotypes) [62]. Lkt is encoded in an operon consisting of four genes *(lktCABD*). *lktA* encodes the protoxin, *lktC* encodes the acylating enzyme essential for post-translational modification and activation of the protoxin, and the products of *lktB* and *lktD* are involved in secretion of the toxin [63,64]. *lktCABD* genes were identified and the predicted protein sequences for the Lkt operon genes from G1 and G2 were aligned (Figure 5). The predicted proteins LktC, LktB and LktD were 95.2%, 99.6% and 100% identical, respectively, suggesting these proteins are under conservation pressure, likely due to their roles in regulating the secretion and function of Lkt [64,65]. In contrast, the predited LktA protein sequences were only 89% identical. Differences in the predicted LktA proteins of G1 and G2 were concentrated in the N-terminal and C-terminal regions of the protein (Figure 5). In leukotoxins the N-terminus is involved in pore-forming activities while the C-terminus is involved in export and host immune system recognition [64]. As such any differences in the N-terminal and/or C-terminal regions may potentially influence the activities of the Lkt and thus may result in virulence differences between the two genotypes.

**Figure 5.**
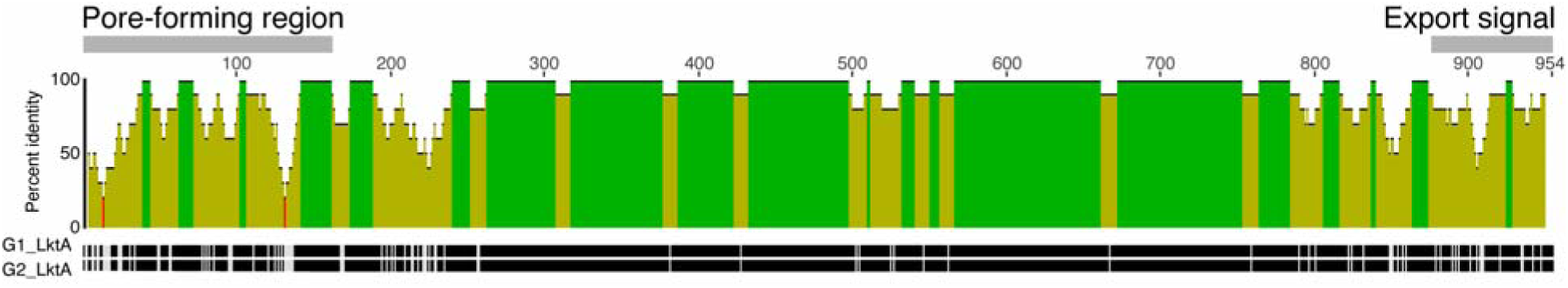
Sequence alignment of G1 and G2 LktA amino acid sequences. Percent sequence identities were calculated with a sliding window size of 10. A larger peak indicates a greater number of matching amino acids within the window for that position. Locations of pore forming domain and export signal domain are indicated above the identity plot. Schematic below the identity plot shows identical (black) and non-identical (light grey) residues in the alignment.

Sequence diversity in LktA within *M. haemolytica* has recently been reported to be greater than within other species in the genus [66]. Davies *et al*. examined the sequence diversity of leukotoxin in 31 *M. haemolytica,* 6 *Mannheimia glucosida,* and 4 *Pasteurella* [*Bibersteinia*] *trehalosi* isolates identified eight *M. haemolytica* allelic variants of *lktA* [63]. Five of the alleles (A6-A10) were identified solely in ovine isolates, allele A3 was only identified in bovine isolates, while A1 and A2 alleles were identified in both ovine and bovine isolates. The A1 allele was subdivided into 5 groups (A1.1-A1.5) with A1.1 solely occurring in bovine isolates from serotype 1 (n = 3) and 6 (n = 1) isolates (corresponding to G2). In bovine isolates from serotype 2 (corresponding to G1), the authors reported detection of alleles A2, A2.1, A2.2, and A3. LktA sequences from genomes in our current study were compared with representative amino acid sequences for each allele and subgroup downloaded from NCBI Genbank. Within G2, 141/146 LktA sequences were 100% identical to the A1.1 allele and another 4/146 were >99.5% identical to A1.1, consistent with the observations of Davies *et al*. (S6-S8 Tables). The only exception was the LktA from Genome 2, that shared 100% amino acid identity with A1.3 and A1.5 (S9 Table).

Interestingly, all but one of the G1 LktA sequences were 99.5-100% identical to the A8.1 allele (Genome 157 was identical to A2) (S6 and S9-10 Tables). The A8.1 allele was found in serotype 2 (G1) isolates by Davies *et al*., however, all the isolates in that study were ovine (n = 3), and A8.1 was also detected in ovine serotype 7 isolates (n = 3). The identification of A8.1 across many bovine origin genomes and nearly complete absence of A2 alleles suggests that G1 from Canada and the U.S.A has diverged from the G1 found in the United Kingdom where most of the isolates used by Davies *et al*. were collected and further demonstrates that A8.1 is not restricted to ovine isolates [63]. Whether there is a relationship between specific LktA alleles and respiratory disease severity in cattle warrants further investigation.

### CAZymes

Carbohydrate active enzymes (CAZymes) are enzymes whose function has been biochemically verified to be involved in the metabolism of carbohydrates. We chose to investigate any differences in CAZymes between the two genotypes since these may indicate differences in ability to metabolize certain carbohydrates, and thus in the nutritional niches of the genotypes [67]. We were particularly interested in enzymes predicted to be secreted into either the periplasmic or extracellular space (i.e. contain a signal peptide) and are thus most likely to be involved in interactions with the external environment. A total of six CAZymes containing signal peptides were identified in the pangenome (S11 Table) (50). At least one CAZyme was predicted in all genomes except for Genome 86/GCF_022404935, which was removed from this analysis. CAZymes identified were members of families GH23 (lytic transglycosylase), GH33 (sialidase), GH13_19 (amylose glucosidase), GT9 (glycosyltransferase), GH102 (lytic transglycosylase) and GH103 (lytic transglycosylase). Only GH103 (homology group 2042729) was genotype associated, present in 60/61 (98.4%) G1 genomes and 2/146 (1.37%) G2 genomes (Table 2). The

GH103 family consists of a group of lytic transglycosylases that cleave the β-1,4 linkage between *N*-acetylmuramoyl and *N*-acetylglucosaminyl residues in peptidoglycan and belong to family 3 of the transglycosylases with structures similar to lysozyme [68,69]. Numerous biological functions have been attributed lytic transglycosylases, however the only clear role is that they are involved in cell wall remodelling [68–71]. Whether this genotype difference in lytic transglycosylases is associated with phenotypic differences in colony morphology of the two genotypes [21] or other differences in cell wall maintenance will require further investigation.

## Conclusions

We identified a total of 422 *Mannheimia haemolytica* genotype differentiating genes (135 G1 specific, 21 G1 associated, 233 G2 specific, and 33 G2 associated), including 125 with unknown functions and 132 that are associated with mobile genetic elements. Genotype associated genes and variants that may contribute to differences in disease association include TonB-dependent transporters, IgA proteases, a highly immunogenic outer membrane protein (PlpE), a lytic transglycosylase, and genotype variants of the leukotoxin gene. The list of genotype differentiating genes provides a shortlist of targets that warrant further investigation to provide a mechanistic understanding of phenotypic differences between the *M. haemolytica* genotypes and inform rational strategies for *M. haemolytica* and bovine respiratory disease diagnostics.

## Acknowledgements

The authors are grateful to Dr. David Palmer (Department of Chemistry, University of Saskatchewan) for assistance with the protein structure alignments, and to Dr. Maarten Voordouw (Department of Veterinary Microbiology, University of Saskatchewan) for assistance with statistical analysis and approaches.

## Supporting Information

**S1 Dataset**. Pangenome database. https://doi.org/10.6084/m9.figshare.28872536.v1

**S1 Table**. Summary of Genomes Included in Pangenome Analysis

**S2 Table**. Accession numbers of sequences used as BLAST queries against pangenome database.

**S3 Table**. Overview of pangenome classification of all homology groups. Genes matching is the number of genes that were found to correspond to the given homology group, as this referred to genes the number may be greater than or less than the number of genomes examined. For pangenome classification: core = group was present in ≥90% of all genomes (187/208); unique = Group was present in exactly one genome; Accessory = Group was present in somewhere between 2-186 genomes. For Genotype classification: Shared = Group was present in ≥90% of the genomes of the given genotype with no consideration of the presence/absence pattern of the other genotype; exclusive = Group was present solely in genomes of the given genotype; Specific = Group was exclusive to one genotype and present in ≥ 90% of that genotype; Associated = Group was present in ≥95% of the genomes of one genotype and >0% and ≤5% of the other genotype.

**S4 Table.** Genotype Specific and Associated Homology Groups. Annotation notes column provides summary of the interpro, pfam and GO functional annotations. Genes matching is the number of genes that were found to correspond to the given homology group, as this referred to genes the number may be greater than or less than the number of genomes examined. For pangenome classification: core = group was present in ≥90% of all genomes (187/208); unique = Group was present in exactly one genome; Accessory = Group was present in somewhere between 2-186 genomes. For phenotype (Genotype) classification: Shared = Group was present in ≥90% of the genomes of the given genotype with no consideration of the presence/absence pattern of the other genotype; exclusive = Group was present solely in genomes of the given genotype; Specific = Group was exclusive to one genotype and present in ≥ 90% of that genotype; Associated = Group was present in ≥95% of the genomes of one genotype and >0% and ≤5% of the other genotype. For manual function categories: Metal Related = Predicted protein function involved in metal metabolism and/or heavy metal resistance, Transporters & Adhesins = Predicted protein is a transporter or adhesin, AMR = Predicted protein is an antimicrobial resistance gene. Regulatory = Predicted protein has a regulatory function, Restriction Endonuclease = Predicted protein is a restriction endonuclease or related to restriction modification, Metabolism = Predicted protein has a function in metabolism such as enzymes involved in carbohydrate degradation, MGE = Predicted proteins are characteristic of mobile genetic elements such as transposases, toxin-antitoxin plasmid maintenance systems, prophage associated proteins etc., Hypothetical = Predicted proteins are hypothetical/putative or a protein with a domain of unknown function (DUF) that lacks any further functional information, Other = Predicted protein could not be categorized into one of the other categories

**S5 Table.** Overview of the number of specific and associated homology groups assigned to each functional category split by genotype classification. Specific = Group was exclusive to the given genotype and present in ≥ 90% of genomes of that genotype, Associated = Group was present in ≥95% of the genomes of one genotype and 0% and ≤5% of the other genotype. For manual function categories: Metal Related = Predicted protein function involved in metal metabolism and/or heavy metal resistance, Transporters & Adhesins = Predicted protein is a transporter or adhesin, AMR = Predicted protein is an antimicrobial resistance gene. Regulatory = Predicted protein has a regulatory function, Restriction Endonuclease = Predicted protein is a restriction endonuclease or related to restriction modification, Metabolism = Predicted protein has a function in metabolism such as enzymes involved in carbohydrate degradation, MGE = Predicted proteins are characteristic of mobile genetic elements such as transposases, toxin-antitoxin plasmid maintenance systems, prophage associated proteins etc., Hypothetical = Predicted proteins are hypothetical/putative or a protein with a domain of unknown function (DUF) that lacks any further functional information, Other = Predicted protein could not be categorized into one of the other categories

**S6 Table**. Pairwise comparisons of amino acid sequences of genotype 1 (G1) and genotype 2 (G2) Leukotoxin A consensus sequences against representatives of LktA alleles from Davies *et al*. 2001

**S7 Table**. Pairwise comparison of all genotype 2 LktA nucleotide sequences.

**S8 Table**. Pairwise comparison of genotype 2 LktA amino acid sequences that had a divergent LktA nucleotide sequence to representative sequences of LktA alleles identified in Davies *et al*. 2001.

**S9 Table**. Pairwise comparison of all genotype 1 LktA nucleotide sequences.

**S10 Table**. Pairwise comparison of genotype 1 LktA amino acid sequences that had a divergent LktA nucleotide sequence to representative sequences of LktA alleles identified in Davies *et al*. 2001.

**S11 Table**. Summary of CAZyme Analysis Results. Only CAZymes that were predicted to contain a signal peptide and were annotated by 3 of the utilized annotation tools (dbCAN_sub, DIAMOND, & HMMER) were included. No CAZymes were predicted in genome 86/GCF_022404935 and so was excluded from this analysis.

**S1 Figure.** Phylogenetic tree of study genomes created using MASH with the ani command in PanTools. G1 and G2 clusters are indicated.

