## Supplementary material for "Identification of genes that differentiate *Mannheimia haemolytica* genotypes 1 and 2 using a pangenome approach": S1 Figure

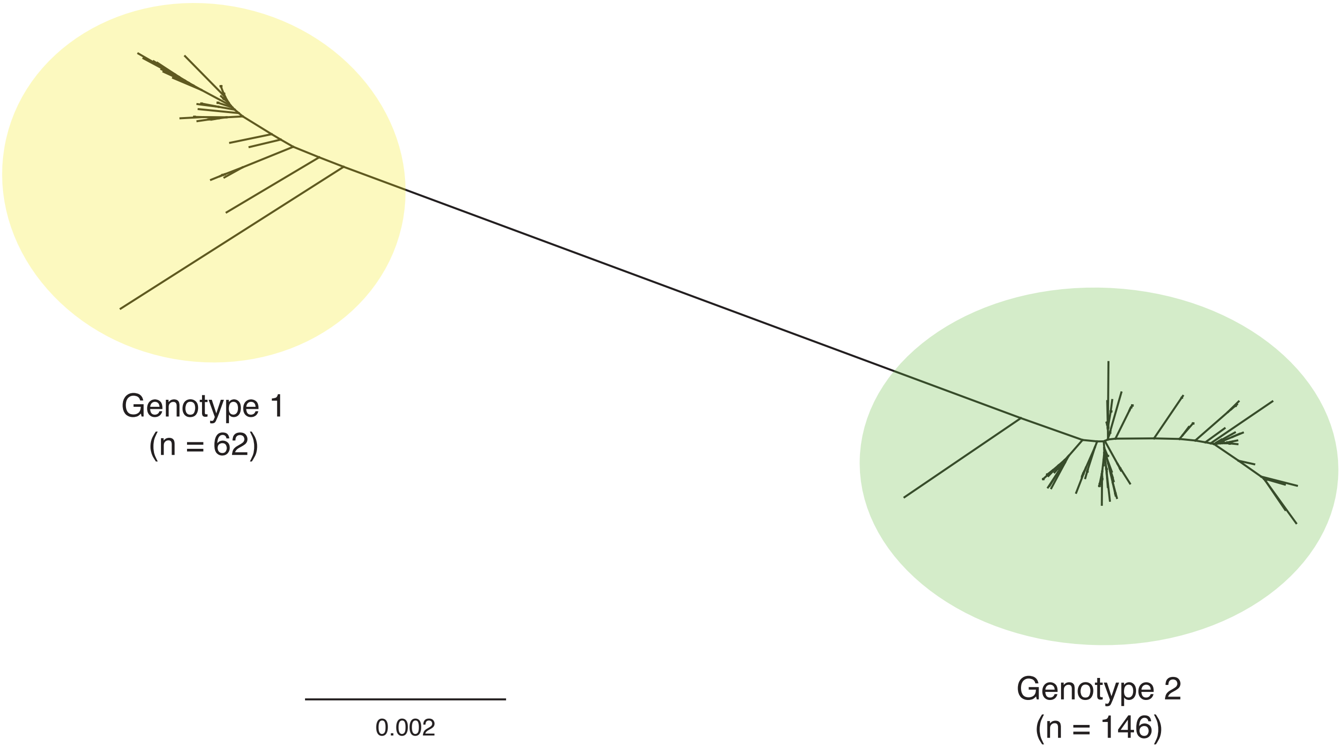


**S1 Figure.** Phylogenetic tree of study genomes created using MASH with the ani command in PanTools. G1 and G2 clusters are indicated.
